# Evolution of drug-binding residues in bacterial ribosomes

**DOI:** 10.1101/2024.10.31.621234

**Authors:** Chinenye L. Ekemezie, Lewis I. Chan, Charlotte R. Brown, Karla Helena-Bueno, Tom A. Williams, Sergey V. Melnikov

## Abstract

Ribosomes from certain bacteria possess divergent drug-binding sites compared to those of *Escherichia coli,* leading to natural evasion or hypersensitivity to antibiotics. However, in the absence of systematic studies, it is unknown whether this observed divergence is a rare exception or a common occurrence among bacterial species. Here, we address this question by reconstructing the evolutionary history of drug-binding residues of the ribosome from the origin of bacteria to the present day. This analysis reveals the extent of natural diversity of ribosomal drug-binding sites between bacterial species, which may inform the development of species-specific antimicrobials and a more accurate and personalized choice of ribosome-targeting drugs for a given pathogen.

## INTRODUCTION

When *Escherichia coli* was introduced as a model for studying bacterial ribosomes, it rapidly became a foundational system for understanding how ribosomes interact with antibiotics (1–4). However, subsequent studies revealed that the sequence of the ribosomal drug-binding residues is not strictly conserved across bacteria, leading to variation in ribosome affinity for antibiotics (5–13). Specifically, species of the human pathogens *Propionibacteria* bear the naturally occurring 16S rRNA substitution G1491U (compared to *E. coli)*, which confers intrinsic resistance to aminoglycosides (12). Some species of *Mycoplasma* carry the 23S rRNA substitution A2057G, which confers resistance to macrolides (13). And the bacterium *Thermus thermophilus* carries the substitutions U1782C/U2586C in the 23S rRNA, rendering this species resistant to tetracenomycin X (8). Without systematic studies, it is currently unknown whether this observed variation in the ribosomal drug-binding sites is rare or common among bacteria.

This lack of systematic studies was caused by two main issues. First, there is a data availability bias, as the majority of genomic sequences for bacterial species belong to only two phyla, Proteobacteria and Firmicutes (14). Consequently, Proteobacteria may be represented by over 10, 000 genome sequences for a single species, while other phyla have only about 20 sequences combined for all species. This disparity skews average conservation scores, favouring rRNA conservation in Proteobacteria and Firmicutes over all bacteria.

The second, more challenging issue is the occurrence of false-positive changes in biological sequences. Previously, datasets of biological sequences were shown to bear spurious mutations resulting from errors of sequencing, assembly or annotation of sequencing data (15, 16). Particularly, since many biological sequences are assembled from multiple shorter reads, public repositories were estimated to contain up to 20% chimeric sequences in which different parts of a given sequence originate from different species instead of a single species (15, 16). Other common errors include sequencing errors, contamination with viral DNA and plasmids, and misannotation of species (15, 16). Consequently, many bacterial species exhibit hundreds of alternative sequence variants of a single gene, complicating the distinction between conserved changes fixed in a lineage from rare mutations and sequence errors.

We addressed these challenges by developing an approach that simplifies the analysis of large biological sequence datasets based on the phylogenetic proximity of bacterial species. This allowed us to assess natural variation at the 82 individual ribosomal drug-binding residues across 8, 809 representative species from all bacterial phyla, providing the first comprehensive atlas of the evolutionary diversity of these medically relevant residues in the active sites of bacterial ribosomes.

## RESULTS

### Evolution-based filtering reveals common variations in ribosomal drug-binding sites

To assess the conservation of drug-binding residues in bacterial ribosomes, we used the public repository of rRNA sequences known as the SILVA dataset NR99 v138.1 because this is the most complete dataset of non-redundant rRNA sequences. This dataset comprises 510, 508 sequences for 16S rRNA and 95, 286 sequences for 23S rRNA (17). We first simplified the dataset by discarding all rRNA sequences of unclear evolutionary origin, and then excluded sequences that originate from phages, plasmids, and eukaryotic hosts of pathogenic bacteria, as well as those with truncations in ribosomal active centres (see **Methods)**. However, even the simplified dataset contained up to 800 alternative rRNA sequences for individual species (**SI_Data 2, 3)**. This illustrated a common obstacle in analysing the evolutionary variations across large groups of species: for example, for *E. coli* alone, we observed 39 dissimilar variants of the ribosomal drug-binding residues, which was partly due to the presence of sequences from clinical strains (**Fig S1a)**.

Therefore, we further simplified the dataset based on the evolutionary proximity of species in the tree of life. In this approach, we selected one representative rRNA sequence per species, choosing the sequences that are most similar to sequences from the closest relatives in the tree (**Fig S1b)**. This approach discarded sequence variants that were specific to a single sequence or strain of a given species but minimized the risk of detecting false positives and thereby revealing the most conserved sequence variants in each bacterial genus (**Fig S1b)**. Using this approach, we identified the most common variants of 82 drug-binding residues in rRNA residues in 8, 809 bacterial species and traced the evolutionary origin and age of these variants (see **Table S1, Fig 1)**. We have summarized our findings in (**SI_Data 4, 5)** to provide the most complete atlas of the conservation of ribosomal drug-binding sites across bacteria.

**Figure 1.**
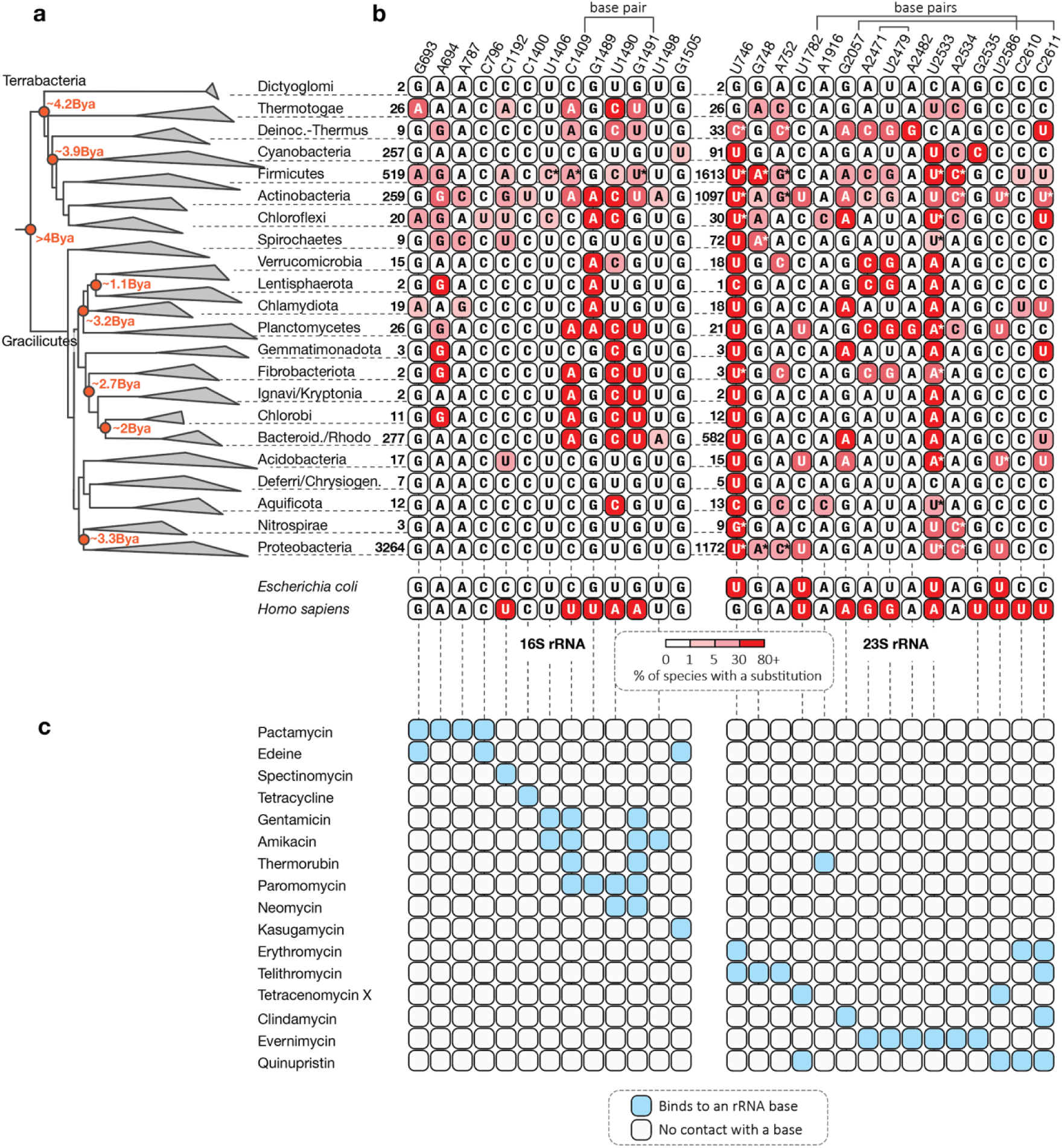
Natural variation in ribosomal drug-binding sites across bacterial phyla. (**a**) The bacterial tree of life calculated based on the conservation of the 16S rRNA illustrates the evolutionary diversification of ribosomal drug-binding residues (**b**) and shows which ribosome-targeting drugs bind to specific residues in rRNA (**c**). Numbers in (**b)** correspond to the total count of representative rRNA sequences that were used to assess substitution profiles of drug-binding residues in rRNA. For simplicity, (**b, c)** display only those rRNA residues that form direct contacts between ribosome-targeting antibiotics and their variable bases (as opposed to the invariant backbone). This panel provides only a simplified summary of variations, while more comprehensive information regarding 8,809 bacterial species can be found in (**SI_Data S4, S5)**. Asterisks indicate sites that have multiple types of substitutions (e.g. G to U and A instead of G to U). Overall, the figure shows that most bacterial phyla exhibit multiple substitutions in their ribosomal drug-binding residues compared to E. coli and to one another. Some of these substitutions likely occurred at least 3 billion years ago (Bya), indicating their ancient origin and widespread distribution across bacteria.

### Phyla-specific variation in ribosomal drug-binding sites

To assess the evolutionary origin and age of substitutions in ribosomal drug-binding sites, we mapped these substitutions onto the bacterial tree of life (**Fig 1)**. This mapping revealed that bacteria likely began to diversify their ribosomal drug-binding residues before their separation into the modern phyla. The earliest substitution likely occurred at least 3.8 billion years ago (Bya) when the branches of Dictyoglomi and Thermotogae separated from other bacteria, bearing the macrolide-binding residue G746 in the 23S rRNA, while the remaining bacteria (specifically of Terrabacteria) bore U746. Subsequently, bacterial ribosomes continued to diversify their drug-binding residues by accumulating lineage-specific substitutions in 28 rRNA bases that directly bind with ribosome-targeting drugs. As a result, we showed that each bacterial phylum has evolved an idiosyncratic set of variations in their ribosomal drug-binding residues (**Fig 1)**.

Compared to *E. coli,* the most dissimilar drug-binding residues we observed among Firmicutes and Actinobacteria. Within these phyla, many species have acquired more substitutions compared to *E. coli* than *E. coli* is to humans. This can be seen, for example, in the order Propionibacteriales, which includes causative agents of skin infections (**SI_Data 5, 6**). Compared to *E. coli,* these species exhibit up to 15 substitutions in their drug-binding residues in the binding sites for pactamycin, neomycin and thermorubin.

Furthermore, some bacteria have substitutions in rRNA bases that are conserved between *E. coli* and humans and therefore viewed as targets of universal inhibitors of protein synthesis. For example, the 16S rRNA residue A694 binds the antibiotic pactamycin and conserved between *E. coli* and humans, which mediates comparable toxicity of pactamycin to both model bacteria and humans and limits pactamycin clinical applications (18, 19). But the laboratory-engineered substitution of this single residue from A to G confers pactamycin resistance (20). Our analysis showed that the A694G substitution is also widespread in nature, specifically among Firmicutes, Actinobacteria, Chloroflexi, Chlamydiota and all members of Spirochaetes (**Fig 1)**. Aside from this substitution, we observed substitutions at 15 drug-binding residues with the same nucleotide state in *E. coli* and humans but different states in other bacteria (**Fig 1)**. Overall, we found that bacteria have been diversifying their ribosomal drug-binding residues since the emergence of the first superphyla, leading to phyla-specific variants of drug-binding sites in bacterial ribosomes.

### Even closely related bacteria can have dissimilar ribosomal drug-binding sites

We found that, after the separation into phyla, bacteria continued to diversify their ribosomal drug-binding residues, resulting in additional substitutions in smaller clades (**Fig 1, SI_Data 4, 5)**. As might be expected, the highest number of these additional substitutions were found in pathogenic bacteria with small genomes, particularly *Rickettsia, Helicobacter, Bartonella, Treponema* and *Mycoplasma* (**Fig 2, SI_Data 4, 5)**. Previously, these species were shown to accumulate genetic changes and evolve more rapidly than non-parasitic bacteria due to weaker natural selection associated with parasitic lifestyles (21–23). Here, we found that this rapid evolution likely extends to the ribosomal drug-binding residues, leading to significant differences even for species even within the same genus. For example, *Mycoplasma* species contain up to 10 substitutions of drug-binding residues compared to each other, leading to a greater divergence between some of these species than they are to *E. coli* (**Fig 2)**.

**Figure 2.**
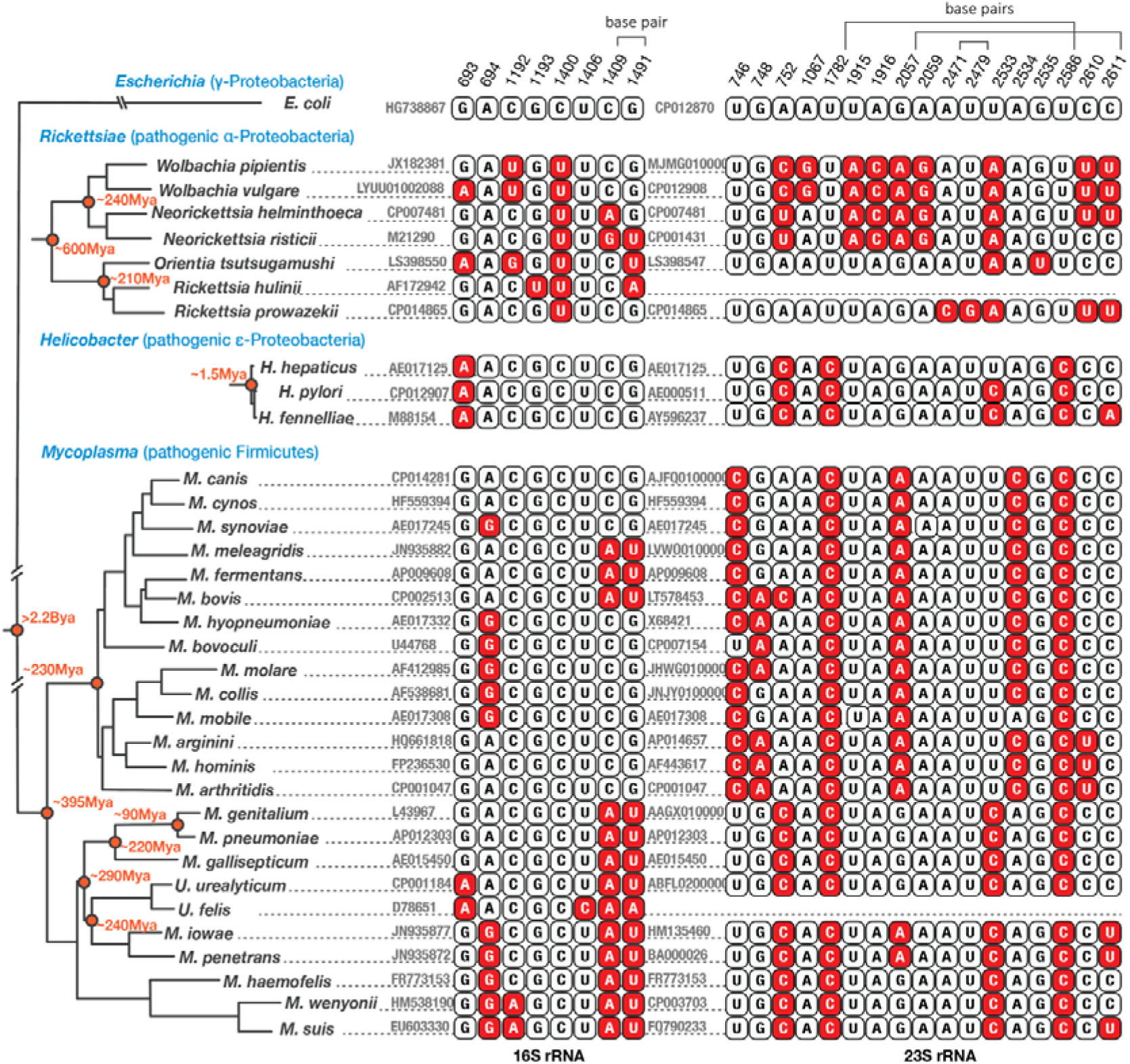
Closely related bacteria can have highly divergent ribosomal drug-binding sites. A phylogenetic tree and multiple sequence alignment illustrate the natural variations in ribosomal drug-binding residues of pathogenic bacteria Mycoplasma, Helicobacter, and Rickettsiae. The estimated ages of bacterial lineages in millions of years (Mya) are derived from genomic and fossil analyses of the corresponding branches (see **Methods)**. Asterisks denote two Mycoplasma species (M. hominis and M. pneumoniae) where the natural divergence of rRNA residues G/A2057 was previously shown to confer intrinsic antibiotic resistance, as observed in an ∼18,000x higher tolerance to 14- and 15-membered macrolides (13). Overall, this analysis reveals that drug-binding sites can diverge rapidly among even closely related bacteria.

Importantly, we found that *Mycoplasma* and other parasitic bacteria tend to bear substitutions which disrupt canonical pairing of the G2057-C2611 and other drug-binding base pairs. For example, in *Rickettsia and Neorickettsia* species, the aminoglycoside-binding base pair C1409-G1491 shows substitutions to A-G, G-U, or C-A pairs, suggesting substantial structural changes in this drug-binding site. Furthermore, rRNA substitutions also involve non-paired residues, such as the pactamycin-binding residues G693 and A694 in the 16S rRNA or the ketolide-binding residues U746 and G748 in the 23S rRNA, among others (**Fig 2**). These examples illustrate that species do not necessarily need to be distantly related to display highly divergent ribosomal drug-binding sites.

### Variants described as resistance-conferring in model bacteria are widespread in nature

We next determined the most common variations in drug-binding residues in bacterial ribosomes. To lower the risk of bias toward overrepresented phyla (9, 24, 18, 25), we first separated rRNA sequences into different phylogenetic groups and then determined representative profiles of individual drug-binding residues for each phylum (**Figs 1; SI_Data 4, 5)**. When rRNAs from different phyla were compared in this way that treats species and phyla more evenly, they revealed high overall variability of ribosomal drug-binding residues in bacteria (**Fig 3, 4)**.

**Figure 3.**
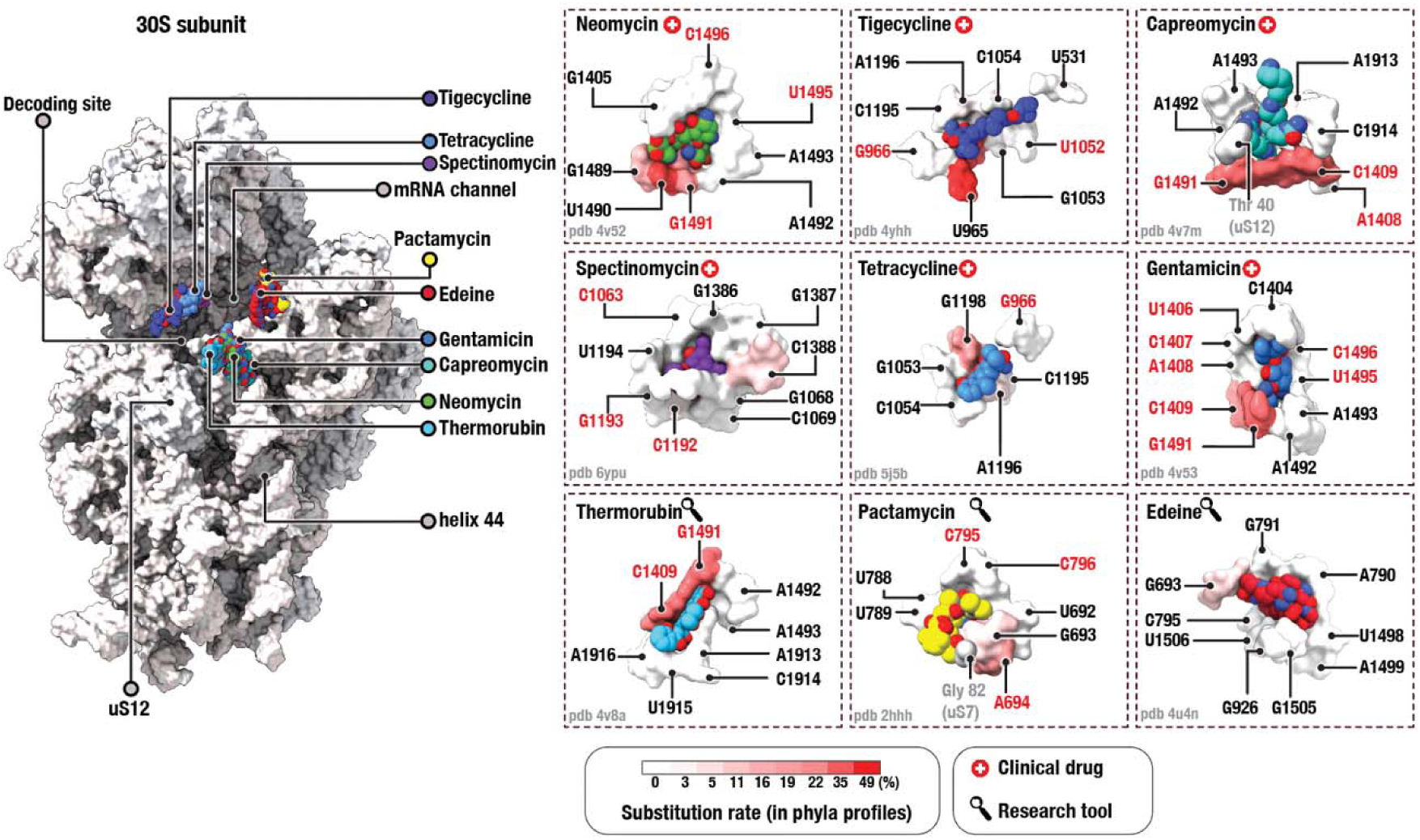
Variable drug-binding sites of the small ribosomal subunit. Structure of the small ribosomal subunit (on the left) and 9 representative drug-binding pockets (on the right) illustrate the conservation of individual drug-binding residues in the 16S rRNA across bacterial phyla. Ribosome-targeting drugs are shown as blobs coloured by atom type, and drug-binding residues—as surfaces coloured by conservation across phyla. Red labels (e.g., C1494 in the neomycin panel) indicate rRNA residues where mutations have been previously shown to confer antibiotic resistance in model bacteria (as summarized in **Table S5)**.

**Figure 4.**
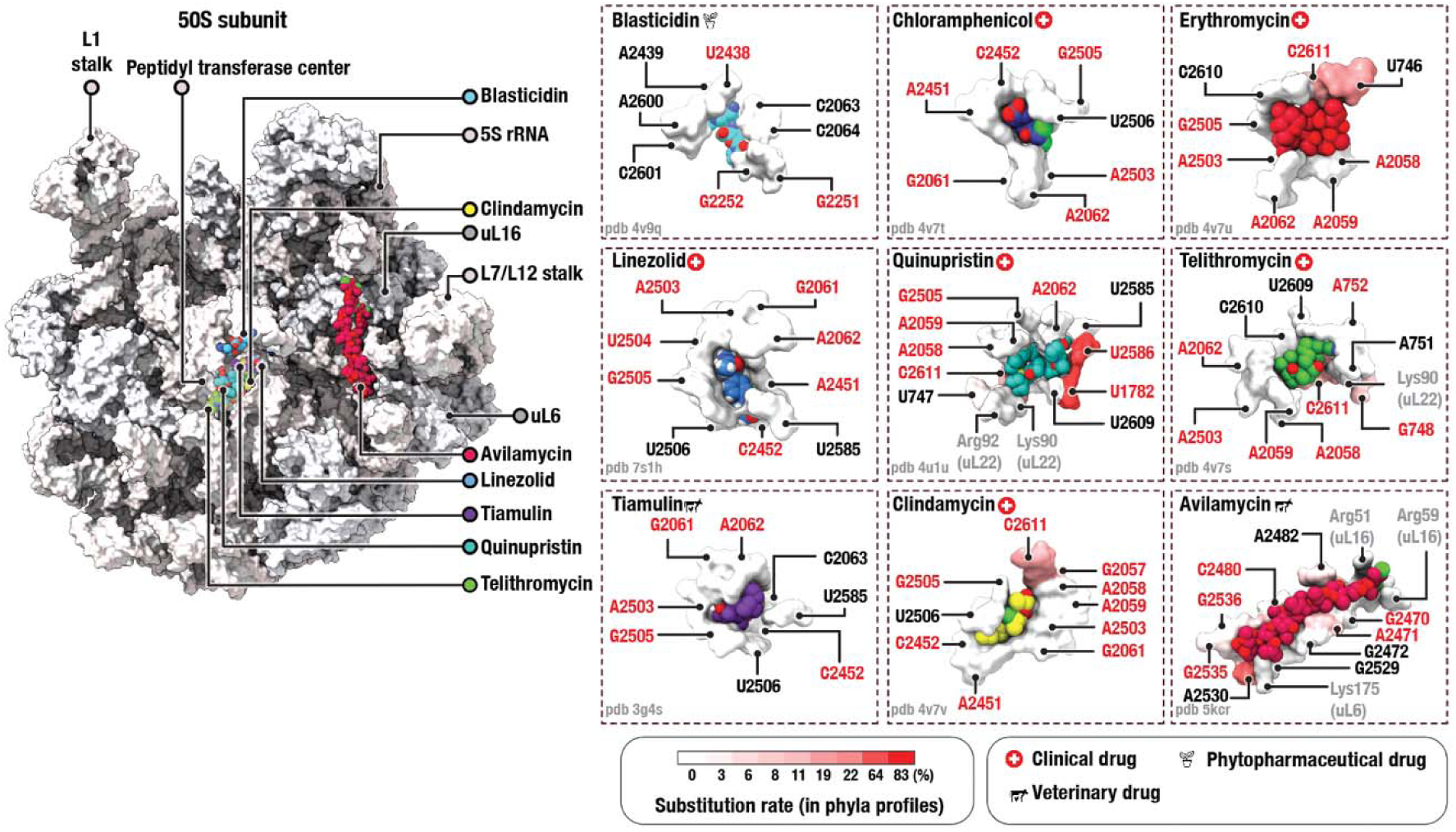
Variable drug-binding sites of the large ribosomal subunit. Structure of the large ribosomal subunit (on the left) and 9 representative drug-binding pockets (on the right) illustrate the conservation of individual drug-binding residues in the 23S rRNA across bacterial phyla. Ribosome-targeting drugs are shown as blobs coloured by atom type, and drug-binding residues—as surfaces coloured by conservation across phyla. Red labels (e.g., U2438 in the blasticidin panel) indicate rRNA residues where mutations have been previously shown to confer antibiotic resistance in model bacteria (as summarized in **Table S6)**.

We finally asked whether the naturally occurring variants identified in our analysis have been previously characterized as resistance-conferring in model bacteria. Our literature review revealed that most natural variants identified in our study were previously described as resistance-conferring in model bacteria, where they led to a 10- to 80, 000-fold higher tolerance to the corresponding ribosome-targeting drug (**Tables S5, S6)**. For example, the 16S rRNA residue C1192, which base pairs with G1064, is recognized by the antibiotic spectinomycin (26). Previous studies showed that spectinomycin binding strictly requires the presence of G at position 1064 and C at position 1192 because mutating either of these residues results in up to a 1, 000x higher spectinomycin resistance (27–29). These high levels of resistance were observed regardless of whether the substitutions maintained base pairing (e.g., C1064-G1192 or A1064-U1192) or not (e.g., U1064-U1192 or G1064-G1192) (27–29).

Furthermore, the effect of these substitutions was observed in multiple model bacteria— including *E. coli, Pasteurella multocida, Salmonella enterica,* and *Borrelia burgdorferi* —where the substitutions C1192U or C1192G resulted in comparable levels of spectinomycin resistance (27–29).

However, our analysis revealed that the C1192U and C1192A variants are not restricted to the previously obtained laboratory strains but frequently occur in nature. Particularly, they are common for most members of the Firmicutes, Actinobacteria, and Chloroflexi phyla, suggesting intrinsic spectinomycin resistance in these species (**Fig 1, SI_Data 4, 5)**. Aside from these C1192 variations, we found that natural variations in 24 drug-binding residues have been characterized as resistance-conferring in model bacterial species (**Tables S5, S6)**. Thus, we found that variants previously described as resistance-conferring in model bacteria are widespread in nature.

## DISCUSSION

### Bacterial phyla have diverged ribosomal drug-binding sites

Prior to our study, the conservation of rRNA residues was estimated using one of the following two approaches. In the first, conservation estimates were derived from sequence alignments, where rRNA sequences were used to calculate the overall conservation of rRNA nucleotides without separation of sequences into phyla and adjustments for sample size of individual clades (7, 9, 11, 24, 25, 30, 31). These analyses were crucial for identifying immutable rRNA bases, but they were biased toward common species and phyla, particularly Proteobacteria, which caused an overestimation of ribosomal drug-binding site conservation. Alternatively, a more accurate and targeted approach involved determining the variations of rRNA drug-binding residues by comparing ribosome structures (6, 7, 18, 19, 32). These analyses provided detailed information about the diversity of ribosomal drug-binding residues, but for only a handful of model organisms.

Here, we described the evolutionary variants of individual rRNA drug-binding residues across more than 8, 000 representative bacteria. We traced the evolutionary history of individual drug-binding residues from the last bacterial common ancestor to the present day, enabling us to estimate variations not only in currently known species but also in those yet to be discovered, based on their position on the tree of life. This analysis revealed that bacterial phyla exhibit a greater diversity of ribosomal drug-binding residues than previously appreciated, illustrating that the occasional findings of divergent variants in species like *Propionibacterium*, *Thermus,* or *Mycoplasma* are a common occurrence in bacterial species.

One implication of our work is that individual bacterial organisms may not serve as reliable generalized models for bacterial ribosomes in the absence of evolutionary analyses. For example, we show that the popular model organism *E. coli* exhibits relatively atypical drug-binding sites for the antibiotics quinupristin, tetracenomycin X, and evernimycin due to the 23S rRNA variations (U1782, U2533, and U2586) because these variants are found in Proteobacteria, but not in most other bacteria. This finding shows that our commonly used division of ribosomes into bacterial and eukaryotic types in the structural studies of ribosomes (6, 18, 33) is incomplete and oversimplified, given the substantial diversity of ribosomal drug-binding residues within the bacterial domain of life.

### Implications for clinical practice

Previously, the discovery of intrinsic antibiotic resistance in *Propionibacteria* and *Mycoplasma* has changed the clinical targeting of these human pathogens (10, 12, 13, 34). For example, *M. pneumoniae,* a respiratory mycoplasma, which has the *E. coli* -type macrolide-binding base G2057 in the 23S rRNA, can be effectively treated with macrolides, lincosamides, streptogramins, and ketolides. However, *M. hominis,* a genital mycoplasma, which bears the A2057G substitution, cannot be treated with 14- and 15-membered macrolides and ketolides due to its natural resistance. For example, this species is 5, 000 more tolerant to the macrolide erythromycin. Therefore, this species of *Mycoplasma* is treated with other antibiotics that bind less divergent sites (13, 34). Thus, the knowledge on structural diversity of ribosomal drug-binding sites resulted in a more accurate selection of ribosome-targeting antibiotics for a given pathogen.

We anticipate a similar revision for a much broader range of bacterial species because our work showed that approximately one in ten representative bacteria carry rRNA substitutions that have been observed in clinical isolates of drug-resistant human pathogens (**SI_Data 4, 5)**. This finding suggests that the intrinsic resistance previously observed in *Mycoplasma, Propionibacterium* and *Thermus* species is likely prevalent among non-model bacteria due to the presence of comparably or even more divergent drug-binding sites. However, our evolutionary analyses have one caveat worth considering: the phenotypic impact of the observed changes at drug-binding sites might, in some cases, be compensated by changes at other sites close in the ribosome structure; thus, the precise effects on drug resistance of the lineage-specific variants we identified will require further study for individual bacteria. Nonetheless, our work presents the first comprehensive overview of the structural variability of ribosomal drug-binding sites in bacteria which may inform a more accurate and personalized choice of ribosome-targeting drugs for a given pathogen.

## MATERIAL AND METHODS

### Annotating ribosomal drug-binding residues

To annotate the drug-binding residues of the ribosome, we used the Protein Data Bank (35) to retrieve previously determined ribosome structures in complex with 35 clinically relevant drugs (26, 36–59), which corresponded to 21 chemical families of ribosome-targeting molecules (**Table S1)**. We then used PyMOL (version 2.5.0) (60) to select ribosomal drug-binding residues located within hydrogen bond distance from the drug, with a distance cut-off of 3.3 Å for structures containing hydrogen atoms and 4.3 Å for structures lacking hydrogen atoms. To identify ribosomal residues that bind ribosome-targeting drugs in a sequence-dependent manner, we then excluded residues that bind drugs with either their sugar moiety or phosphate backbone of rRNA nucleotides. This allowed us to select only those rRNA residues that bind drugs using their nucleotide bases (**Table S2)**.

### Creating a high-fidelity dataset of rRNA sequences for evolutionary analyses

To enable accurate analysis of rRNA sequences, we have used the public repository of rRNA sequences, SILVA, particularly the dataset NR99 v138.1 that contained 510, 508 non-redundant 16S rRNA sequences corresponding to 40, 239 species (17, 609 genera) (“SSU dataset”) and 95, 286 23S rRNA sequences from 14, 849 species (7, 837 genera) (“LSU dataset”) from all domains of life (17). To filter out false-positive sequence variations, we have devised a 10-step data reduction strategy to bypass the errors in sequence datasets that stem from automated annotation of sequencing data and sequencing mistakes, such as chimeric sequences (where a single sequence is assembled from fragments from different species), sequences with misannotated phylogeny, or sequences from non-genomic DNA and pseudogenes (**Table S3 and S4)**. In this reduction strategy, the initial SILVA datasets were first reduced by eliminating sequences from non-bacterial species as well as from unidentified bacteria for which we could not confirm the identity of an organism. This included sequences whose headers contained the words “unidentified”, “uncultured”, “metagenome”, “cluster” and “sp.”, as well as sequences whose names began with a lowercase letter. The obtained datasets were then further reduced by eliminating rRNA sequences that are encoded by plasmids and viruses. We then removed sequences of apparently poor sequencing quality, based on the presence of ‘ambiguous’ nucleotides represented by symbols such as “R”, “X”, “N”, and “S” instead of the standard set of RNA bases “A”, “G”, “C”, and “U”. This resulted in the 23S rRNA dataset containing 56, 154 sequences from 5, 726 bacterial species (from 1, 678 genera) and the 16S rRNA dataset containing 94, 486 sequences from 14, 132 of bacterial species (from 2, 967 genera) (**SI_Data S2 and S3)**.

### Assessing of drug-binding residues in bacterial rRNA

To enable the analysis of rRNA conservation across bacterial species, the reduced sequence libraries of 16S and 23S rRNA were used for multiple sequence alignments. To reduce gaps and alignment errors, the datasets were first separated into sequences from individual bacterial phyla, and then the multiple sequence were produced using MAFFT v7.4908 (61) with default settings (FFT-NS-2) for each phylum. To assess the conservation of drug-binding residues, the aligned sequences of 16S and 23S rRNA were reduced to the sequences of drug-binding residues only (**SI_Data S4, S5)**. To remove potential “false-positive” sequence variations that originate from truncations, the errors of sequencing, alignment or genome assembly, we removed sequences that contained truncations in the ribosomal drug-binding sites. The resulting dataset was further reduced to a single sequence per organism using the following rule: whenever the same organism contained several sequence variants, we used only one variant, which was identical (or the most similar) to the sequences of the adjacent members on the tree of life. This allowed us to identify mutations that were a common characteristic of a clade of species and eliminate sequencing and annotation artifacts, as well as mutations that were characteristic of an individual strain (**SI_Data S4, S5)**.

### Mapping the conservation of bacterial ribosomal drug-binding residues on ribosome structure

To highlight conservation of the drug-binding residues in the structure of the bacterial ribosome, we used ChimeraX v1.6.19. First, we generated the profile sequence of the ribosomal drug-binding residues for each bacterial phylum in our dataset (shown in **SI_Data 5, 6)**. We then aligned the profile sequences and calculated the percentage of conservation for each drug-binding residue in the aligned phyla-specific profile sequences. The resulting scores were then shown in (Figure 2).

### Assessing the conservation of drug-binding residues in rRNA operons of the same species

To assess potential variability of rRNA drug-binding residues between rRNA operons from the same species, we have repeated our analysis of rRNA-binding residues by using the manually curated dataset of rRNA operons rrnDB (version 5.8) (62) instead of the SILVA dataset. Our analysis showed that the drug-binding residues of rRNA encoded by different operons of the same species remain conserved, thereby excluding the possible variability of antimicrobial resistance between rRNA produced by different operons of the same bacterial species.

### Tracing the evolutionary origin of variations in bacterial ribosomal drug-binding sites

To trace the evolutionary history of variations in the ribosomal drug-binding residues, we used 16S rRNA sequences listed in (**SI_Data 5, 6)**, we used the dated bacterial tree generated using Mega11 with default settings (63), rooting the tree based on recent analyses (64, 65). The relative age of each mutation was estimated based on a published bacterial time tree (65) or the original studies of *Mycoplasma* (66, 67), *Helicobacter* (21, 68) and *Rickettsiales* (69).

## DATA AVAILABILITY

Data accompanying this work were deposited in FigShare: https://doi.org/10.6084/m9.figshare.27370026.v1

## AUTHOR CONTRIBUTIONS

C.L.E., L.I.C. and S.V.M. conceived the study and performed the analysis; C.R.B. assisted with the automation. K.H.B. contributed to the analysis of rRNA variations in species with small genomes. All authors wrote the manuscript.

## ACKNOWLEDGEMENTS

We thank members of Melnikov lab for their help in the preparation of this manuscript and Heath Murray, Katarzyna Mickiewicz, Robert Hirt and Claudia Schneider (Newcastle University, UK) for their critical comments.

