## Supplementary Information for "Evolution of drug-binding residues in bacterial ribosomes"

**Supplementary Figures**

Figure S1 | Evolution-based filtering reveals common natural variants of the drug-binding residues of bacterial ribosomes.

Figure S2 | Common model bacteria bear many variable drug-binding residues.

**Supplementary Tables**

Table S1 | Ribosome-targeting antibiotics analysed in this study.

Table S2 | Ribosomal drug-binding residues assessed in this study.

Table S3 | Quality control steps to prepare 16S rRNA sequences for the analysis.

Table S4 | Quality control steps to prepare 23S rRNA sequences for the analysis.

Table S5 | 16S rRNA mutations known to confer drug resistance.

Table S6 | 23S rRNA mutations known to confer drug resistance.

**Supplementary Data (**DOI: <10.6084/m9.figshare.27370026>)

- To download the data: <https://figshare.com/ndownloader/files/50103723>

Supplementary Data 1 | Bacteria analysed in this study.

Supplementary Data 2 | Sequences 16S rRNA used in this study.

Supplementary Data 3 | Sequences 23S rRNA used in this study.

Supplementary Data 4 | Aligned 16S rRNA drug-binding residues.

Supplementary Data 5 | Aligned 23S rRNA drug-binding residues.

Supplementary Data 6 | Scripts used in this study.

**Supplementary Figures**

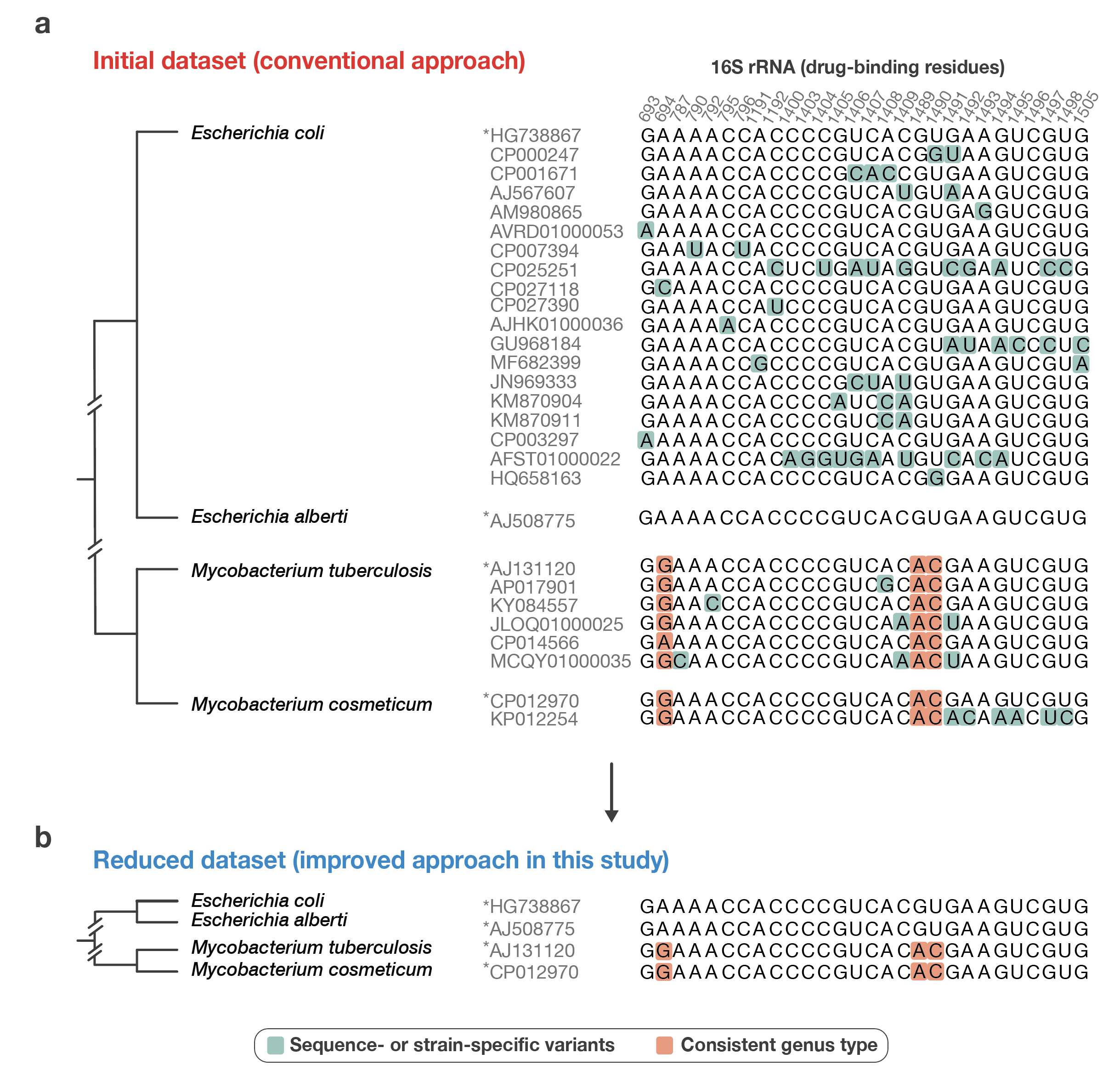

**Figure S1 | Evolution-based filtering reveals common natural variants of the drug-binding residues of bacterial ribosomes.** (**a**) A fragment of aligned 16S rRNA sequences from the SILVA dataset (NR99 v138.1) illustrates the apparent variability of drug-binding residues in bacterial ribosomes among representative members of *Escherichia* and *Mycobacterium* species. Notably, the extensive number of sequences in current biological repositories suggests that *E. coli* alone contains at least 19 dissimilar variants of drug-binding residues. This high apparent degree of variability within a single organism complicates the identification of common variations in drug-binding residues and raises questions about the authenticity of these variants—specifically, how many of them are sequencing errors or strain-specific variants and how many are fixed in species or larger bacterial clades. (**b**) A reduced alignment of rRNA sequences demonstrates our methodology in which we represent each species by a single rRNA sequence, by filtering out the variants that are not shared (at least) between sister rRNA sequences on the tree of life. This approach markedly simplifies the analysis of large libraries of rRNA sequences. Although this method omits variations that are specific to individual sequences or individual bacterial strains, it effectively minimizes stochastic errors, making it possible to identify natural variations that are shared by bacterial genera or broader clades. We have applied this approach to assess conservation of drug-binding residues in 510,508 rRNA sequences from 8,809 bacterial species.

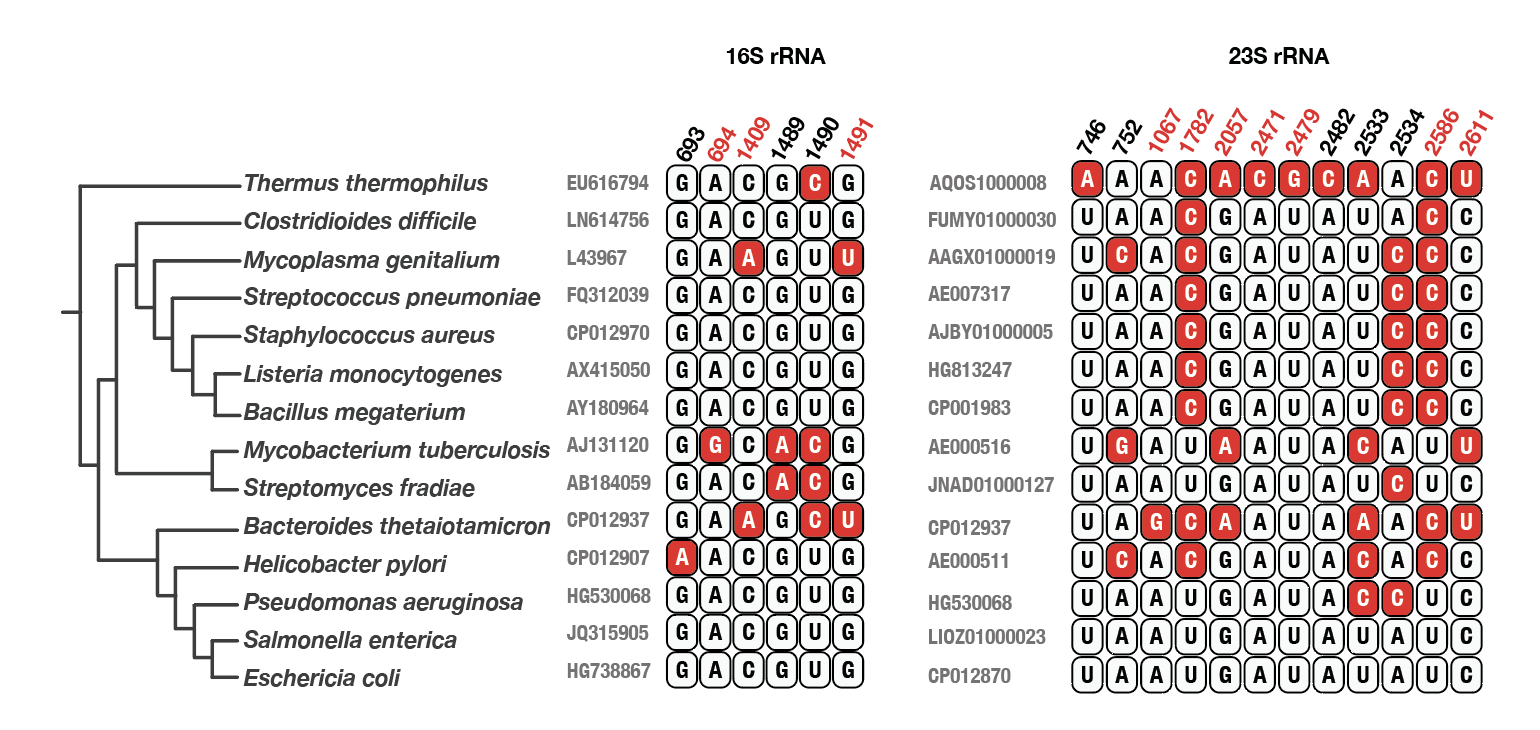

**Figure S2 | Common model bacteria bear many variable drug-binding residues.** Ribosome structures show the position of ribosomal drug-binding sites and highlight the location of variable rRNA drug-binding residues among some common model bacterial species (these rRNA residues are highlighted in red). Residue numbers highlighted in red indicate rRNA bases where mutations have been shown to confer drug resistance. The panel illustrates that certain drug-binding residues of the ribosome are not conserved between many commonly studied bacteria.

**Table S1 | Ribosome-targeting antibiotics which binding sites are assessed in this study.**

| Drug | Family | Subunit | Binding site | Organism | PDP ID | Application | Origin | Year |
| --- | --- | --- | --- | --- | --- | --- | --- | --- |
| Linezolid | Oxazolidinone | 50S | PTC | *E. coli* | 7S1H | Clinical | Synthetic | 2021 |
| Erythromycin | Macrolide | 50S | Tunnel | *E. coli* | 4V7U | Clinical | Natural | 2014 |
| Clindamycin | Lincosamide | 50S | PTC/  Tunnel | *E. coli* | 4V7V | Clinical | Semi-synthetic | 2014 |
| Telithromycin | Ketolide | 50S | Tunnel | *E. coli* | 4V7S | Clinical | Semi-synthetic | 2014 |
| Chloramphenicol | Amphenicol | 50S | PTC | *E. coli* | 4V7T | Clinical | Synthetic | 2014 |
| Avilamycin | Orthosomycin | 50S | A-site | *E. coli* | 5KCR | Veterinary | Natural | 2016 |
| Evernimycin | Orthosomycin | 50S | A-site | *E. coli* | 5KCS | Clinical | Natural | 2016 |
| Quinupristin | Streptogramin B | 50S | Tunnel | *E. coli* | 4U1U | Clinical | Semi-synthetic | 2014 |
| Dalfopristin | Streptogramin A | 50S | PTC | *E. coli* | 4U24 | Clinical | Semi-synthetic | 2014 |
| Tiamulin | Pleuromutilin | 50S | PTC | *H. marismortui* | 3G4S | Veterinary | Semi-synthetic | 2009 |
| Retapamulin | Pleuromutilin | 50S | PTC | *D. radiodurans* | 2OGO | Clinical | Semi-synthetic | 2007 |
| Virginiamycin | Streptogramin | 50S | PTC | *E. coli* | 4U25 | Veterinary | Natural | 2014 |
| Puromycin | Aminonucleoside | 50S | PTC | *H. marismortui* | 1Q81 | Research | Natural | 2000 |
| Sparsomycin | Sparsomycin | 50S | PTC | *H. marismortui* | 1VQ8 | Antitumor, Antibacterial | Natural | 2005 |
| Carbomycin | Macrolide | 50S | PTC/  Tunnel | *H. marismortui* | 1K8A | Veterinary | Natural | 2001 |
| Azithromycin | Macrolide | 50S | Tunnel | *H. marismortui* | 1M1K | Clinical | Semi-synthetic | 2002 |
| Tetracenomycin X | Tetracenomycin | 50S | Tunnel | *E. coli* | 6Y69 | Research, universal | Natural | 2020 |
| Thiostrepton | Thiopeptides | 50S | GTP-ase site | *D. radiodurans* | 3CF5 | Veterinary | Natural | 2008 |
| Blasticidin S | Peptidyl nucleoside | 50S | P-site | *T. thermophilus* | 4V9Q | Phytophar-maceutical | Natural | 2014 |
| Kasugamycin | Aminoglycoside | 30S | E-site | *T. thermophilus* | 2HHH | Research | Natural | 2006 |
| Spectinomycin | Aminocyclitol aminoglycoside | 30S | Neck | *E. coli* | 4V56 | Clinical | Natural | 2014 |
| Neomycin | Aminoglycoside | 30S | A-site | *E. coli* | 4V52 | Clinical | Natural | 2014 |
| Tigecycline | Glycylcycline | 30S | A-site | *T. thermophilus* | 4YHH | Clinical | Semi-synthetic | 2015 |
| Tetracycline | Tetracycline | 30S | A-site | *E. coli* | 5J5B | Clinical | Natural | 2016 |
| Streptomycin | Aminoglycoside | 30S | A-site | *T. thermophilus* | 4DR3 | Clinical | Natural | 2012 |
| Edeine | Peptide | 30S | E-site | *S. cerevisiae* | 4U4N | Research | Natural | 2014 |
| Pactamycin | Aminoglycoside | 30S | E-site | *T. thermophilus* | 4W2G | Universal inhibitor | Natural | 2014 |
| Thermorubin | Anthraceno-pyranone | 30/50S | A-site | *T. thermophilus* | 4V8A | Research tool | Natural | 2012 |
| Negamycin | Pseudodipeptide | 30S | A-site | *E. coli* | 4WF1 |  | Natural | 2002 |
| Capreomycin | Tuberactinomycin | 30S | A-site | *T. thermophilus* | 4V7M | Clinical | Natural | 2010 |
| Viomycin | Tuberactinomycin | 30S | A-site | *T. thermophilus* | 6LKQ | Clinical | Natural | 2020 |
| Hygromycin B | Aminoglycoside | 30S | A-site | *E. coli* | 4V64 | Clinical | Natural | 2008 |
| Paromomycin | Aminoglycoside | 30S | A-site | *L. donovani* | 6AZ1 | Clinical | Natural | 2017 |
| Gentamicin | Aminoglycoside | 30S | A-site | *E. coli* | 4V53 | Clinical | Natural | 2007 |
| Amikacin | Aminoglycoside | 30S | A-site | *A. baumanii* | 6YPU | Clinical | Semi-synthetic | 2020 |

**Table S2 | rRNA residues that directly contact ribosome-targeting drugs using their aromatic bases.**

| 16S rRNA residue | Antibiotics | Position in the alignment  (SI_Data 4) |
| --- | --- | --- |
| G693 | Edeine, and Pactamycin | 1 |
| A694 | Pactamycin | 2 |
| A787 | Pactamycin | 3 |
| A790 | Edeine | 4 |
| G791 | Edeine | 5 |
| A792 | Edeine, Kasugamycin | 6 |
| A794 | Edeine, Kasugamycin | 7 |
| C795 | Edeine, Kasugamycin, Pactamycin | 8 |
| C796 | Pactamycin | 9 |
| G926 | Edeine, Kasugamycin | 10 |
| C1054 | Negamycin, Tetracycline, Tigecycline | 11 |
| C1063 | Spectinomycin | 12 |
| G1064 | Spectinomycin | 13 |
| C1066 | Spectinomycin | 14 |
| A1191 | Spectinomycin | 15 |
| C1192 | Spectinomycin | 16 |
| G1193 | Spectinomycin | 17 |
| C1400 | Tetracycline | 18 |
| C1403 | Hygromycin B, | 19 |
| C1404 | Amikacin, Gentamicin, Hygromycin B, Viomycin, | 20 |
| G1405 | Amikacin, Gentamicin, Hygromycin B, Neomycin | 21 |
| U1406 | Amikacin, Gentamicin, Hygromycin B | 22 |
| C1407 | Amikacin, Gentamicin, Neomycin, Paromomycin, Viomycin | 23 |
| A1408 | Amikacin, Capreomycin, Gentamicin, Neomycin, Paromomycin, Thermorubin, Viomycin | 24 |
| C1409 | Amikacin, Capreomycin, Gentamicin, Paromomycin, Thermorubin | 25 |
| G1489 | Paromomycin | 26 |
| U1490 | Neomycin, Paromomycin | 27 |
| G1491 | Amikacin, Capreomycin, Gentamicin, Neomycin, Paromomycin, Thermorubin, Viomycin | 28 |
| A1492 | Amikacin, Capreomycin, Viomycin | 29 |
| A1493 | Amikacin, Capreomycin, Hygromycin B, and Viomycin | 30 |
| G1494 | Amikacin, Capreomycin, Gentamicin, Hygromycin B, Neomycin, Paromomycin, Viomycin | 31 |
| U1495 | Amikacin, Gentamicin, Hygromycin B, Neomycin, Paromomycin, Viomycin | 32 |
| C1496 | Amikacin, Gentamicin, Hygromycin B, Neomycin, Viomycin | 33 |
| G1497 | Amikacin, Hygromycin B | 34 |
| U1498 | Amikacin | 35 |
| G1505 | Edeine, Kasugamycin | 36 |
| 23S rRNA residues | **Antibiotics** | **Position in the alignment**  **(SI_Data 5)** |
| U746 | Azithromycin, Erythromycin, and Telithromycin | 1 |
| G748 | Telithromycin | 2 |
| A752 | Telithromycin | 3 |
| A1067 | Thiostrepton | 4 |
| A1095 | Thiostrepton | 5 |
| A1096 | Thiostrepton | 6 |
| U1782 | Tetracenomycin X, Quinupristin | 7 |
| A1913 | Capreomycin, Viomycin, and Thermorubin | 8 |
| C1914 | Capreomycin, Viomycin | 9 |
| U1915 | Thermorubin | 10 |
| A1916 | Thermorubin | 11 |
| G2057 | Clindamycin | 12 |
| A2058 | Azithromycin, Carbomycin, Clindamycin, Erythromycin, Quinupristin, and Telithromycin | 13 |
| A2059 | Azithromycin, Carbomycin, Clindamycin, Erythromycin, Quinupristin, and Telithromycin, | 14 |
| G2061 | Azithromycin, Carbomycin, Clindamycin, Dalfopristin, Linezolid, Puromycin, Retapamulin, Tiamulin, and Virginiamycin | 15 |
| A2062 | Carbomycin, Chloramphenicol, Erthromycin, Tetracenomycin X, Dalfopristin, Linezolid, Quinupristin, Retapamulin, Sparsomycin, Telithromycin, Tiamulin, and Virginiamycin | 16 |
| C2063 | Dalfopristin, Puromycin, Retapamulin, and Virginiamycin | 17 |
| G2251 | Blasticidin | 18 |
| G2252 | Blasticidin | 19 |
| A2439 | Blasticidin, and Virginiamycin | 20 |
| A2451 | Carbomycin, Chloramphenicol, Clindamycin, Dalfopristin, Linezolid, Puromycin, Retapamulin, Sparsomycin, Tiamulin, and Virginiamycin | 21 |
| C2452 | Carbomycin, Chloramphenicol, Clindamycin, Dalfopristin, Puromycin, Retapamulin, Sparsomycin, Tiamulin, and Virginiamycin | 22 |
| A2469 | Avilamycin, and Evernimycin | 23 |
| G2470 | Avilamycin, and Evernimycin | 24 |
| A2471 | Avilamycin, and Evernimycin | 25 |
| A2478 | Avilamycin, and Evernimycin | 26 |
| U2479 | Evernimycin | 27 |
| C2480 | Evernimycin | 28 |
| A2482 | Evernimycin | 29 |
| A2503 | Azithromycin, Carbomycin, Chloramphenicol, Clindamycin, Erythromycin, Dalfopristin, Linezolid, Retapamulin, Telithromycin, Tiamulin, and Virginiamycin | 30 |
| U2504 | Chloramphenicol, Clindamycin, Dalfopristin, Linezolid, Retapamulin, and Virginiamycin | 31 |
| G2505 | Azithromycin, Clindamycin, Erythromycin, Quinupristin, and Retapamulin | 32 |
| U2506 | Chloramphenicol, Clindamycin, Dalfopristin, Puromycin, and Retapamulin | 33 |
| U2533 | Evernimycin | 34 |
| A2534 | Evernimycin | 35 |
| G2535 | Avilamycin, and Evernimycin | 36 |
| G2553 | Puromycin | 37 |
| G2583 | Dalfopristin, and Puromycin | 38 |
| U2584 | Dalfopristin, Puromycin, and Sparsomycin | 39 |
| U2585 | Tetracenomycin X, Linezolid, Puromycin, Tiamulin, and Virginiamycin | 40 |
| U2586 | Tetracenomycin X, and Quinupristin | 41 |
| A2587 | Tetracenomycin X | 42 |
| A2602 | Puromycin | 43 |
| U2609 | Tetracenomycin X, Quinupristin, and Telithromycin | 44 |
| C2610 | Erythromycin, and Quinupristin | 45 |
| C2611 | Azithromycin, Carbomycin, Clindamycin, Erythromycin, Quinupristin, and Telithromycin | 46 |

**Table S3 | Steps of quality control to simplify the 16S rRNA SILVA dataset.**

| Quality control steps | Criteria | Search words | Filtered records | Sequences left | Unique species | Unique genus |
| --- | --- | --- | --- | --- | --- | --- |
| Silva raw data |  |  | 0 | 510,508 | 40,578 | 17,731 |
| Remove non genomic sequences | Plasmid or phage encoded sequence | Phage, phage, prophage, Plasmid, plasmid | 184 | 510,324 | 40,577 | 17,731 |
| Remove organellar encoded sequence | Sequence from mitochondrial or chloroplast origin | Mitochondria and chloroplast | 6,237 | 504,087 | 39,761 | 17,401 |
| Remove metagenomic sequence |  | metagenome | 8,792 | 495,295 | 39,761 | 17,401 |
| Remove poorly characterized sequences | Sequences of unknown taxonomy, unknown species and unknown genus (small caps genus name) | aff., sp., archaeon, associated, bacterium, Candidatus, cf., clone, cluster, culture, environmental, Epixenosomes, eukaryotum, euryarchaeote, -group, Incertae Sedis, isolate, -like, phytoplasma, proteobacterium, snow, symbiont, Thermogales str, unclassified, unidentified, uncultured, unknown family | 356,249 | 139,046 | 32,631 | 14,201 |
| Remove low quality sequence | Nucleotides different from the canonical A, G, C and U | Nucleotides designated as: B, D, H, K, M, N, R, S, V, W, X, Y and Z | 15,857 | 123,189 | 32,631 | 14,201 |
| Remove records from archaeal and eukaryotic origin | Records annotated as archaea and eukaryotes | Archaea and eukaryote | 28,703 | 94,486 | 14,132 | 2,967 |
| Remove unnatural mutants | Mutant record overlapping with non-mutant record | None | 18,427 | 76,059 | 14,130 | 2,967 |
| Remove truncated active sites | Sequence with deletion in any of their drug-binding residues | Ribosomal drug-binding residues designated as gap (-) | 13,912 | 62,147 | 8,728 | 2,171 |
| Remove duplicates | Species with more than one record | None | 53,194 | 8,953 | 8,728 | 2,171 |
| Remove misclassified records | Eukaryotic species classified as bacteria, bacterial species classified as a bacterial lineage other than their taxonomically characterized lineage and other poorly characterized sequence | None | 146 | 8,809 | 8,671 | 2,126 |

**Table S4 | Steps of quality control to simplify the 23S rRNA SILVA dataset.**

| Quality control steps | Criteria | Search words | Filtered records | Sequences left | Unique species | Unique genus |
| --- | --- | --- | --- | --- | --- | --- |
| Silva LSU raw dataset |  |  | 0 | 95,286 | 14,912 | 7,871 |
| Remove non genomic sequences | Plasmid or phage encoded sequence | Phage, phage, prophage, Plasmid, plasmid | 182 | 95,104 | 14,912 | 7,871 |
| Remove organellar encoded sequence | Sequence from mitochondrial or chloroplast origin | Mitochondria and chloroplast | 2,543 | 92,561 | 14,062 | 7,405 |
| Remove metagenomic sequence |  | metagenome | 1,516 | 91,045 | 14,062 | 7,405 |
| Remove poorly characterized sequences | Sequences of unknown taxonomy, unknown species and unknown genus (small caps genus name) | aff., sp., archaeon, associated, bacterium, Candidatus, cf., clone, cluster, culture, environmental, Epixenosomes, eukaryotum, euryarchaeote, -group, Incertae Sedis, isolate, -like, phytoplasma, proteobacterium, snow, symbiont, Thermogales str, unclassified, unidentified, uncultured, unknown family | 19,929 | 71,116 | 13,661 | 7,134 |
| Remove low quality sequence | Nucleotides different from the canonical A, G, C and U | Nucleotides designated as: B, D, H, K, M, N, R, S, V, W, X, Y and Z | 3,440 | 67,676 | 11,674 | 5,898 |
| Remove records from archaeal and eukaryotic origin | Records annotated as archaea and eukaryotes | Archaea and eukaryotes | 11,522 | 56,154 | 5,726 | 1,678 |
| Remove unnatural mutants | Non conserved record overlapping with conserved record | None | 158 | 55,996 | 5,726 | 1,678 |
| Remove truncated active sites | Sequence with deletion in any of their drug-binding residues | Ribosomal drug-binding residues designated as gap (-) | 9,272 | 46,724 | 4,422 | 1,252 |
| Remove duplicates | Bacterial species with more than one record in our dataset | None | 42,260 | 4,464 | 4,422 | 1,252 |
| Remove misclassified records | Eukaryotic species classified as bacteria, bacterial species classified as a bacterial lineage other than their taxonomically characterized lineage and other poorly characterized sequence | None | 5 | 4,459 | 4,422 | 1,252 |
| Extract Alphaproteobacteria records from the original Silva LSU dataset | | | | | | |
| Extract Alphaproteobacteria record from the original Silva dataset | Remove all records not corresponding to Alphaproteobacteria | Alphaproteobacteria | 90,814 | 4,472 | 1,232 | 588 |
| Remove truncated active sites | Sequence with deletion in any of their drug-binding residues | Ribosomal drug-binding residues designated as gap (-) | 166 | 4,306 | 1,999 | 575 |
| Remove poorly characterized sequences | Sequences of unknown taxonomy, unknown species and unknown genus (small caps genus name) | sp., proteobacterium, metagenome, bacterium, uncultured, Incertae, Candidatus, endosymbionts | 1,799 | 2,507 | 692 | 250 |
| Remove duplicates | Bacterial species with more than one record in our dataset | None | 1,809 | 698 | 692 | 250 |

**Table S5 | 16S rRNA mutations that are known to confer drug resistance.**

| Base | Organism | Organism type | Mutation | Effect | Ref |
| --- | --- | --- | --- | --- | --- |
| A694 | *Halobacterium halobium* | Lab mutant | A to G | 4-fold higher MIC of spectinomycin | (1) |
| C795 | *Halobacterium halobium* | Lab mutant | C to U | 4-fold higher MIC of spectinomycin | (1) |
| C796 | *Halobacterium halobium* | Lab mutant | C to U | 4-fold higher MIC of spectinomycin | (1) |
| C912 | *Nicotiana plumbaginfolia* | Lab mutant | C to U | Streptomycin resistance | (2,3) |
| C912 | *Euglena gracilis* | Lab mutant | C to U | Streptomycin resistance | (4) |
| C912 | *Chlamydomonas reinhardtii* | Wild type | C to U | Streptomycin resistance | (5) |
| C912 | *Nicotiana tabacum* |  | C to U | Streptomycin resistance | (6) |
| C912 | *Nicotiana tabacum* |  | C to A | Streptomycin resistance | (7) |
| C912 | *Escherichia coli* | Lab mutant | C to G | 4-fold increase in resistance to streptomycin | (8) |
| A913 | *Escherichia coli* |  | A to G | Streptomycin resistance | (9) |
| A914 | *Thermus thermophilus* | Lab mutant | A to G | Streptomycin resistance | (10) |
| A914 | *Escherichia coli* |  | A to C | Streptomycin resistance | (11) |
| A915 | *Thermus thermophilus* | Lab mutant | A to G | Streptomycin resistance | (10) |
| U965 | *Escherichia coli* |  | A to G | Streptomycin resistance | (9) |
| U965 | *Helicobacter pylori* | Clinical isolate | A to G | Tetracycline resistance | (12) |
| G966 | *Escherichia coli* | Lab mutant | G to U | 4-fold higher MIC of tetracycline | (13) |
| G966 | *Escherichia coli* | Lab mutant | G to U | 2-fold higher MIC for negamycin | (13) |
| G966 | *Escherichia coli* | Lab mutant | G to U | 4-fold higher MIC for tetracycline and tigecycline | (14) |
| U1052 | *Escherichia coli* | Lab mutant | U to G | 8-fold higher MIC for negamycin | (13) |
| U1052 | *Escherichia coli* | Lab mutant | U to G | 4-fold higher MIC for tetracycline | (15) |
| U1052 | *Escherichia coli* | Lab mutant | U to G | 8-fold higher MIC for negamycin | (15) |
| G1058 | *Escherichia coli* | Lab mutant | G to C | 4-fold higher MIC for negamycin and tetracycline | (13) |
| U1060 | *Escherichia coli* | Lab mutant | U to A | 8-fold higher MIC for negamycin | (13) |
| C1063 | *Escherichia coli* | Lab mutant | C to U | 8-fold higher MIC for spectinomycin | (16) |
| G1064 | *Escherichia coli* | Lab mutant | G to A, G | Spectinomycin resistance | (17) |
| G1064 | *Neisseria meningitidis* & *Neisseria gonorrheae* | Clinical isolate | G to C | Spectinomycin resistance | (18) |
| C1066 | *Escherichia coli* | Lab mutant | C to U | 32- fold higher MIC for spectinomycin | (16) |
| C1066 | *Escherichia coli* | Lab mutant | C to U | 62-fold higher MIC for spectinomycin | (19) |
| A1191 | *Chlamydomonas reinhardtii* |  | A to C, A to G | Streptomycin resistance | (5) |
| A1191 | *Borrelia burgdorferi* | Lab mutant | A to G | >2,200-fold higher MIC for spectinomycin | (20) |
| A1191 | *Chlamydomonas reinhardtii* (chloroplasts) | Lab mutant | A to G | 500-fold higher MIC for spectinomycin | (21) |
| C1192 | *Escherichia coli* | Lab mutant | C to G | 32-fold higher MIC for spectinomycin | (16) |
| C1192 | *Pasteurella multocida* | Veterinary | C to G | 512-fold higher MIC for spectinomycin | (22) |
| C1192 | *Salmonella typhimurium (wild type), E. coli (lab strain)* | Lab mutant | C to U | 500-fold higher MIC for spectinomycin | (19) |
| C1192 | *Borrelia burgdorferi* | Lab mutant | C to U | >2,250-fold higher MIC for spectinomycin | (20) |
| C1192 | *Escherichia coli* | Lab mutant | C to U | Spectinomycin resistance | (23) |
| C1192 | *Mycobacterium smegmatis* | Lab mutant | C to G | Spectinomycin resistance | (24) |
| G1193 | *Escherichia coli* | Lab mutant | G to A | 32-fold higher MIC for spectinomycin | (16) |
| G1193 | *Chlamydia psittaci* rRNA expressed in *E. coli* | Lab mutant | G to C | Spectinomycin resistance | (25) |
| A1197 | *Escherichia coli* | Lab mutant | A to U | 16-fold higher MIC for negamycin | (26) |
| G1386 | *Nicotiana tabacum* | Lab mutant | G to A | Spectinomycin resistance | (27) |
| U1406 | *Mycobacterium smegmatis* | Lab mutant | U to C | 64-fold higher MIC for hygromycin | (28) |
| U1406 | *Thermus thermophilus* | Lab mutant | U to C | 50-fold, 10-fold, 5-fold and 2-fold increase in resistance to kanamycin, gentamicin resistance, hygromycin and capreomycin respectively | (29) |
| U1406 | *Thermus thermophilus* | Lab mutant | U to A | 5-fold, 50-fold and 1,000-fold increase in resistance to hygromycin, kanamycin and gentamicin, respectively | (29) |
| U1406 | *Thermus thermophilus* | Lab mutant | U to G | 10-fold, 10-fold, 5-fold and 2-fold increase in resistance to kanamycin, gentamicin, hygromycin and capreomycin | (29) |
| U1406 | *Escherichia coli* | Lab mutant | U to A | 64-fold higher MIC for gentamicin C and tobramycin, 128-fold higher MIC for kanamycin A and G418 | (30) |
| U1406 | *Mycobacterium smegmatis* | Lab mutant | U to C | 8-fold increase in resistance to neamine and ribostamycin, 16-fold increase in resistance to paromomycin, and 4-8-fold increase in resistance to lividomycin | (31) |
| C1407 | *Escherichia coli* | Lab mutant | C to U | Impairs paromomycin binding to the ribosome | (32) |
| A1408 | *Mycobacterium smegmatis* | Lab mutant | A to G | 64-fold increase in resistance to paromomycin, more than 1,024-fold increase in resistance to neomycin, and more than 1,024 increase in resistance to gentamicin, tobramycin and kanamycin | (33) |
| A1408 | *Borrelia burgdorferi* | Clinical isolate | A to G | 90-fold increase in resistance to kanamycin and more than 240-fold increase in resistance to gentamicin | (20) |
| A1408 | *Thermus thermophilus* | Lab mutant | A to G | 25-fold higher MIC for streptomycin, 250-fold higher MIC for apramycin, 20-fold MIC for paromomycin, 500-fold MIC for neomycin, 1,000-fold MIC for gentamicin, 1,500-fold increase in resistance to kanamycin, and 20-fold increase in resistance to capreomycin | (29) |
| A1408 | *Mycobacterium avium, Mycobacterium abscessus* | Clinical isolates | A to G | 16-fold higher MIC for amikacin compared to the recommended breakpoint of 16 for susceptible isolates | (30) |
| A1408 | *Mycobacterium smegmatis* | Lab mutant | A to G | 16-fold higher MIC for neamine, more than 128 higher MIC for ribostamycin, 512-1,024-fold higher MIC for neomycin, 64-fold higher MIC for paromomycin resistance, and 32-fold higher MIC for lividomycin | (31) |
| C1409 | *Mycobacterium smegmatis* | Lab mutant | C to G | 32-fold higher MIC for paromomycin, 4-fold for neomycin, >1024-fold higher MIC for geneticin, 4-fold higher MIC for gentamicin and 16-fold higher MIC for tobramycin | (36) |
| C1409 | *Mycobacterium smegmatis* | Lab mutant | C to U | 8-fold higher MIC for paromomycin and tobramycin, 128-fold higher MIC for geneticin, 16-fold higher MIC for gentamicin | (36) |
| C1409 | *Mycobacterium smegmatis* | Lab mutant | C to U | 4-8-fold higher MIC for paromomycin, 8-fold higher MIC for gentamicin, 8-16-fold higher MIC for tobramycin and kanamycin. | (33) |
| C1409 | *Thermus thermophilus* | Lab mutant | C to G | 5-fold higher MIC for neomycin, 25-fold higher MIC for streptomycin, 50-fold higher MIC for paromomycin, 100-fold higher MIC for apramycin, 200-fold higher MIC for gentamicin and kanamycin and 20-fold higher MIC for capreomycin | (29) |
| C1409 | *Mycobacterium smegmatis* | Lab mutant | C to U | 4-fold higher MIC for neamine, 8-fold higher MIC for ribostamycin, 8-fold higher MIC for lividomycin and paromomycin | (31) |
| C1409 | *Mycobacterium smegmatis* | Lab mutant | C to G | 2-fold higher MIC for neamine, 16-32-fold higher MIC for paromomycin, 64-fold higher MIC for lividomycin and ribostamycin | (31) |
| G1491 | *Mycobacterium smegmatis* | Lab mutant | G to A | 64-fold higher MIC for paromomycin | (33) |
| G1491 | *Thermus thermophilus* | Lab mutant | G to A | 2-fold higher MIC for kanamycin, 5-fold higher MIC for paromomycin, 20-fold higher MIC for capreomycin and 50-fold higher MIC for apramycin | (29) |
| G1491 | *Mycobacterium smegmatis* | Lab mutant | G to U/C | 512-fold higher MIC for paromomycin | (36) |
| G1491 | *Mycobacterium smegmatis* | Lab mutant | G to A | 2-fold higher MIC for neamine, 4-fold higher MIC for neomycin, 16-fold higher MIC for ribostamycin, 64-fold higher MIC for paromomycin and 256-fold increase higher MIC for lividomycin | (31) |
| G1491 | *Mycobacterium smegmatis* | Lab mutant | G to C | 16-fold higher MIC for neamine, ~32-fold higher MIC for neomycin 128-fold higher MIC for ribostamycin, 512-fold higher MIC for paromomycin | (31) |
| G1491 | *Mycobacterium smegmatis* | Lab mutant | G to U | 16-fold higher MIC for neomycin, 16-fold higher MIC for neamine, 128-fold higher MIC for ribostamycin, 512-fold higher MIC for paromomycin and lividomycin | (31) |
| U1495 | *Thermus thermophilus* | Lab mutant | U to C | 2-fold higher MIC for kanamycin and 5-fold higher MIC for hygromycin | (29) |
| U1495 | *Mycobacterium smegmatis* | Lab mutant | U to A | 8-fold higher MIC for neomycin, 16-fold higher MIC for neamine 128-fold higher MIC for ribostamycin, 512-fold higher MIC for paromomycin and lividomycin | (31) |
| U1495 | *Mycobacterium smegmatis* | Lab mutant | U to C | 8-fold higher MIC for neamine, 16-fold higher MIC for ribostamycin, 8-fold higher MIC for neomycin, 128-fold higher MIC for paromomycin and 64-fold higher MIC for lividomycin | (31) |
| C1496 | *Mycobacterium smegmatis* | Lab mutant | C to U | 32-fold higher MIC for hygromycin | (28) |
| C1496 | *Mycobacterium avium* | Clinical isolates | C to U | 8-fold higher MIC for amikacin | (34,35) |
| U1498 | *Mycobacterium smegmatis* | Lab mutant | U to C | 16-fold higher MIC for hygromycin | (28) |
| U1498 | *Mycobacterium abscessus* | Clinical isolates | U to A | 16-fold higher MIC for amikacin | (34,35) |

**Table S6 | 23S rRNA mutations that are known to confer drug resistance.**

| Base | Organism | Organism type | Mutation | Effect | Reference |
| --- | --- | --- | --- | --- | --- |
| G748 | *Mycoplasma bovis* | Lab mutant | G to A | 4-fold higher MIC for lincomycin, 16-fold higher MIC for tilmicosin, and 64-fold higher MIC for tylosin | (37) |
| G748 | *Mycoplasma bovis* | Veterinary isolate | G to A | 1,024-fold higher MIC for tylosin, tilomicin, gamithromycin, tildipirosin | (38) |
| A1067 | *Escherichia coli* | Lab mutant | A to C, A to U | Reduced ribosome affinity to thiostrepton to about 35% of the wild-type strain | (39) |
| A1095 | *Escherichia coli* | Lab mutant | A to C, A to U | Significantly decreased thiostrepton affinity in ribosomes bearing this mutation | (40) |
| U1782 | *Escherichia coli* | Lab mutant | U to C | 16-fold higher MIC for tetracenomycin X | (41) |
| G2057 | *Propionibacterium acnes* | Clinical isolate | G to A | Erythromycin resistance | (42) |
| G2057 | *Mycoplasma fermentans,* | Type strain | G to A | >18,000-fold higher MIC for erythromycin, >2,000-fold higher MIC for clarithromycin, >130-fold higher MIC for azithromycin, 64-fold higher MIC for quinupristin, 35-fold higher MIC to telithromycin, 8-fold higher MIC for josamycin, tylosin, and 4-fold higher MIC for midecamycin, pristinamycin and spiramycin, | (43) |
| G2057 | *Mycoplasma pulmonis* | Type strain | G to A | >18,000-fold higher MIC for erythromycin, >2,000-fold higher MIC for clarithromycin, >530-fold higher MIC for azithromycin, >133-fold higher MIC for midecamycin >66-fold higher MIC for josamycin, 35-fold higher MIC for telithromycin, 32-fold higher MIC for spiramycin, 4-fold higher MIC for pristiniamycin | (43) |
| G2057 | *Escherichia coli* | Lab mutant | G to A | Chloramphenicol and erythromycin resistance | (44) |
| A2058 | *Propionibacterium acnes* | Clinical isolate | A to G | Erythromycin, tylosin, spiramycin, josamycin, and clindamycin resistance | (42) |
| A2058 | *Streptococcus pneumoniae* | Clinical isolate | A to G | 512-fold higher MIC for clarithromcyin, 256-fold higher MIC for azithromycin, 64-fold higher MIC for clindamycin, 32-fold higher MIC for midekamycin | (45) |
| A2058 | *Escherichia coli* | Lab mutant | A to G | >6-fold higher MIC for erythromycin, 7.5-fold higher MIC for clarithromycin | (46) |
| A2058 | *Treponema denticola* | Lab mutant | A to G | Erythromycin resistance | (47) |
| A2058 | *Mycobacterium smegmatis* | Lab mutant | A to G | 512-fold higher MIC for telithromycin, 32-fold higher MIC for carbomycin, 8-fold higher MIC for spiramycin, josamycin and desmycosin, 2-fold higher MIC for tylosin | (48) |
| A2058 | *Mycobacterium smegmatis* | Lab mutant | A to C | 4,096-fold higher MIC for telithromycin, 256-fold higher MIC for carbomycin, and josamycin, 32-fold higher MIC for spiramycin, 16-fold higher MIC for desmycosin, 4-fold higher MIC for tylosin | (48) |
| A2058 | *Mycoplasma pneumoniae* | Clinical isolate | A to G | 32,000-fold higher MIC for azithromycin, 17,000-fold higher MIC for clarithromycin and erythromycin, 260-fold higher MIC for midecamycin, 130-fold higher MIC for josamycin, 64-fold higher MIC for clindamycin, 32-fold higher MIC for lincomycin | (49) |
| A2058 | *Mycoplasma pneumoniae* | Clinical isolate | A to C | 17,000-fold higher MIC for erythromycin and clarithromycin, 8,000-fold higher MIC for azithromycin, 1,000-fold higher MIC for midecamycin, 500-fold higher MIC for josamycin, 66-fold higher MIC for rokitamycin, 8-fold higher MIC for lincomycin and clindamycin | (49) |
| A2058 | *Streptococcus pneumoniae (originally clinical isolate)* | Lab mutant | A to G | 2,000-fold higher MIC for azithromycin, erythromycin and clarithromycin, 125-fold higher MIC for lincomycin, 100-fold higher MIC for telithromycin, 31-fold higher MIC for clindamycin, 16-fold higher MIC for spiramycin, 4-fold higher MIC for streptogramin B | (50) |
| A2058 | *Mycoplasma bovis* | Lab mutant | A to U | 64-fold higher MIC for tylosin, 16-fold higher MIC for tilmicosin, 4-fold higher MIC for lincomycin | (37) |
| A2058 | *Mycobacterium smegmatis* | Lab mutant | A to G | 64-fold higher MIC for clindamycin, 2-fold higher MIC for valnemulin | (51) |
| A2058 | *Mycobacterium smegmatis* | Lab mutant | A to G | 128-fold higher MIC for clindamycin, 64-fold higher MIC for erythromycin and azithromycin, 8-fold higher MIC for spiramycin, and josamycin, 2-fold higher MIC for tylosin | (52) |
| A2058 | *Escherichia coli* | Lab mutant | A to U | Oleandomycin, niddamycin, tylosin, spiramycin, lincomycin, clindamycin and osteogrycin B resistance | (53) |
| A2509 | *Mycobacterium smegmatis* | Lab mutant | A to G | 4,096-fold higher MIC for telithromycin, 256-fold higher MIC for carbomycin and josamycin, 64-fold higher MIC for tylosin and desmycosin, 32-fold higher MIC for spiramycin | (48) |
| A2059 | *Streptococcus pneumoniae (originally clinical isolate)* | Lab mutant | A to G | 20,000-fold higher MIC for azithromycin, 1,000-fold higher MIC for spiramycin, 625-fold higher MIC for erythromycin, 312-fold higher MIC for clairthromycin, 10-fold higher MIC for clindamycin, 7.8-fold higher MIC for lincomycin, 3-fold higher MIC for telithromycin, | (50) |
| A2059 | *Mycoplasma pneumoniae* | Clinical isolate | A to G | 32,000-fold higher MIC for azithromycin, 17,000-fold higher MIC for erythromycin, 2,000-fold higher MIC for josamycin, 4,266-fold higher MIC for midecamycin, 533-fold higher MIC for rikotamycin, 8-fold higher MIC for lincomycin, and clindamycin | (49) |
| A2059 | *Propionibacterium acnes* | Clinical isolate | A to G | Erythromycin, tylosin, spiramycin, josamycin, clindamycin, azithromycin, and pristinamycin resistance | (42) |
| G2061 | *Escherichia coli* | Lab mutant | G to C | This mutation is deleterious to *E. coli* cells | (54) |
| G2061 | *Thermus thermophilus* | Lab mutant | G to A | Tiamulin and chloramphenicol resistance | (55) |
| G2061 | *Thermus thermophilus* | Lab mutant | G to U | Tiamulin resistance | (55) |
| A2062 | *Mycoplasma bovis* | Lab mutant | A to U, A to C | 64-fold higher MIC for tylosin, 16-fold higher MIC for tilmicosin | (37) |
| A2062 | *Mycobacterium hominis* | Type culture | A to G, A to U | 2,133-fold higher MIC for josamycin, and miocamycin | (56) |
| A2062 | *Deinococcus radiodurans* | Lab mutant | A to C | Linezolid and pleuromutilin resistance | (57) |
| A2062 | *Halobacterium halobium* | Lab mutant | A to C | 1,024-fold higher MIC for spiramycin, 64-fold higher MIC for josamycin, and tylosin, 32-fold higher MIC for pristinamycin I, 2-fold higher MIC for pristinamycin II | (58) |
| A2062 | *Halobacterium halobium* | Lab mutant | A to C | 26-fold higher MIC for lincomycin | (59) |
| G2252 | *Thermus aquaticus* | Lab mutant | G to A | Caused drastic reduction in peptidyl transferase activity | (60) |
| U2438 | *Halobacterium halobium* | Lab mutant | U to C | Amicetin resistance | (61) |
| A2451 | *Thermus thermophilus* | Lab mutant | A to U | Tiamulin and chloramphenicol cross-resistance | (55) |
| C2452 | *Halobacterium halobium* | Lab mutant | C to U | 53-fold higher MIC for lincomycin | (59) |
| C2452 | *Thermus thermophilus* | Lab mutant | C to U | Tiamulin resistance | (55) |
| C2452 | *Deinococcus radiodurans* | Lab mutant | C to U | Linezolid, chloramphenicol and anisomycin cross-resistance | (57) |
| C2452 | *Haloarcula marismortui* | Lab mutant | C to U | 50-fold higher MIC for anisomycin | (62) |
| C2452 | *Sulfolobus acidocaldarius* | Lab mutant | C to U | Celesticetin, chloramphenicol and carbomycin resistance | (63) |
| A2453 | *Halobacterium halobium* | Lab mutant | A to C | 5-fold higher MIC for lincomycin | (59) |
| A2453 | *Halobacterium halobium* | Lab mutant | A to G | 43-fold higher MIC for lincomycin | (59) |
| A2469 | *Streptococcus pneumoniae* | Lab mutant | A to C | 16-fold higher MIC for avilamycin | (64) |
| G2470 | *Halobacterium halobium* | Lab mutant | G to U | 220-fold higher MIC for avilamycin | (65) |
| A2471 | *Halobacterium halobium* | Lab mutant | A to G | 2,000-fold higher MIC for evernimycin | (66) |
| A2471 | *Halobacterium halobium* | Lab mutant | A to C | 5,400-fold higher MIC for evernimycin | (66) |
| A2471 | *Halobacterium halobium* | Lab mutant | A to G, A to C | 220-fold higher MIC for avilamycin | (65) |
| A2478 | *Halobacterium halobium* | Lab mutant | A to C | 220-fold higher MIC for avilamycin | (65) |
| A2478 | *Halobacterium halobium* | Lab mutant | A to C | 5,400-fold higher MIC for evernimycin | (66) |
| U2479 | *Halobacterium halobium* | Lab mutant | U to C | 220-fold higher MIC for avilamycin | (65) |
| U2479 | *Halobacterium halobium* | Lab mutant | U to C | 5,400-fold higher MIC for evernimycin | (66) |
| C2480 | *Halobacterium halobium* | Lab mutant | C to U | 220-fold higher MIC for avilamycin | (65) |
| C2480 | *Halobacterium halobium* | Lab mutant | C to A | 5,400-fold higher MIC for evernimycin | (66) |
| C2480 | *Halobacterium halobium* | Lab mutant | C to U | 5,400-fold higher MIC for evernimycin | (66) |
| C2480 | *Streptococcus pneumoniae* | Lab mutant | C to U | 16-fold higher MIC for evernimycin | (64) |
| C2499 | *Halobacterium halobium* | Lab mutant | C to U | 10-fold higher MIC for lincomycin | (59) |
| U2500 | *Halobacterium halobium* | Lab mutant | U to C | 66-fold higher MIC for lincomycin | (59) |
| U2500 | *Thermus thermophilus* | Lab mutant | U to A | Tiamulin resistance | (55) |
| A2503 | *Mycoplasma gallisepticum* | Lab mutant | A to U | 128-fold higher MIC for erythromycin, 64-fold higher MIC for chloramphenicol, 62.5-fold higher MIC for valnemulin, 32-fold higher MIC for florfenicol, 8-fold higher MIC for lincomycin | (52) |
| A2503 | *Mycobacterium smegmatis* | Lab mutant | Ato G | 4-fold higher MIC for linezolid, 4-fold higher MIC for chloramphenicol | (51) |
| A2503 | *Mycobacterium smegmatis* | Lab mutant | A to U | 16-fold higher MIC for josamycin and clindamycin, 8-fold higher MIC for spiramycin and linezolid, 4-fold higher MIC for valnemulin and chloramphenicol | (52) |
| U2504 | *Brachyspira hyodysenteriae, Brachyspira pilosicol* | Lab mutant | U to C | Tiamulin and chloramphenicol resistance | (67) |
| U2504 | *Halobacterium halobium* | Lab mutant | U to G | 60-fold higher MIC for lincomycin | (59) |
| U2504 | *Mycobacterium smegmatis* | Lab mutant | U to G | 16-fold higher MIC for chloramphenicol, 4-fold higher MIC for linezolid, 2-fold higher MIC for valnelium and clindamycin | (51) |
| U2504 | *Mycobacterium smegmatis* | Lab mutant | U to G | 16-fold higher MIC for chloramphenicol and florfenicol, 4-fold higher MIC for spiramycin, linezolid, valnemulin, and josamycin, 2-fold higher MIC for clindamycin | (52) |
| U2504 | *Deinococcus radiodurans* | Lab mutant | U to C | Linezolid resistance | (57) |
| U2504 | *Staphylococcus aureus, Enterococus faecium and coagulase negative Staphylococci* | Clinical isolate | U to A | Linezolid resistance | (68) |
| U2504 | *Escherichia coli* | Lab mutant | U to 𝛹 | 16-fold higher MIC for clindamycin, 8-fold higher MIC for linezolid, 4-fold higher MIC for tiamulin | (69) |
| G2505 | *Mycobacterium smegmatis* | Lab mutant | G to A | 8-fold higher MIC for linezolid, 4-fold higher MIC for chloramphenicol | (51) |
| G2505 | *Deinococcus radiodurans* | Lab mutant | G to A | Linezolid resistance | (57) |
| G2505 | *Enterococcus faecium* | Clinical isolate | G to A | 16-fold higher MIC for linezolid | (70) |
| A2534 | *Staphylococcus epidermidis* | Clinical | C to U | 3-fold higher MIC for linezolid | (71) |
| G2535 | *Streptococcus pneumoniae* | Lab mutant | G to A | 8-fold higher MIC for evernimycin | (64) |
| G2535 | *Halobacterium halobium* | Lab mutant | G to A | 170-fold higher MIC for evernimycin | (66) |
| G2535 | *Halobacterium halobium* | Lab mutant | G to A | 22-fold higher IC_50_ for avilamycin | (65) |
| G2535 | *Enterococcus feacalis* | Veterinary isolate | G to A, G to U | Evernimycin resistance | (72) |
| G2536 | *Streptococcus pneumoniae* | Lab mutant | G to C | 8-fold higher MIC for evernimycin | (64) |
| G2553 | *Escherichia coli* | Lab mutant | G to C | Caused severe growth defects in cells grown on erythromycin containing media | (73) |
| U2586 | *Escherichia coli* | Lab mutant | U to G, U to A | 16-fold higher MIC for tetracenomycin X | (41) |
| U2586 | *Escherichia coli* | Lab mutant | U to C | 8-fold higher MIC for tetracenomycin X | (41) |
| U2609 | *Escherichia coli* | Lab mutant | U to A | 4-fold higher MIC for tetracenomycin X | (41) |
| U2609 | *Escherichia coli* | Lab mutant | U to G | 8-fold higher MIC for tetracenomycin X | (41) |
| C2611 | *Escherichia coli* | Lab mutant | C to U | 5-fold higher MIC for erythromycin and clarithromycin | (46) |
| C2611 | *Mycoplasma pneumoniae* | Clinical isolate | C to G | 500-fold higher MIC for erythromycin, 60-fold higher MIC for clarithromycin, 15-fold higher MIC for azithromycin, 4-fold higher MIC for midecamycin | (49) |
| C2611 | *Streptococcus pneumoniae* | Lab mutant (originally clinical isolate) | C to A | 39-fold higher MIC for clarithromycin, 32-fold higher MIC for streptogramin B, 16-fold higher MIC for azithromycin and erythromycin, 4-fold higher MIC for spiramycin | (50) |
| C2611 | *Streptococcus pneumoniae* | Lab mutant (originally clinical isolate) | C to G | 5,000-fold higher MIC for clarithromycin, 2,000-fold higher MIC for erythromycin, 128-fold higher MIC for streptogramin B, 125-fold higher MIC for azithromycin, 8-fold higher MIC for spiramycin, 4-fold higher MIC for lincomycin | (50) |
| C2611 | *Escherichia coli* | Lab mutant | C to U | Erythromycin resistance | (74) |
